# The effects of explicit and implicit information on modulation of corticospinal excitability during hand-object interactions

**DOI:** 10.1101/2022.03.20.485033

**Authors:** Guy Rens, Marco Davare, Vonne van Polanen

**Affiliations:** The Brain and Mind Institute, University of Western Ontario, London, Ontario N6A 3K7, Canada; KU Leuven, Leuven Brain Institute, 3001 Leuven, Belgium; Department of Clinical and Movement Neurosciences, UCL Queen Square Institute of Neurology, University College London, London, United Kingdom; Faculty of Life Sciences and Medicine, King’s College London, London SE1 1UL, United Kingdom; Movement Control and Neuroplasticity Research Group, Department of Movement Sciences, Biomedical Sciences group, KU Leuven, 3001 Leuven, Belgium

**Author notes:** **Corresponding Author:** Guy Rens, The Brain and Mind Institute, University of Western Ontario, Ontario N6A 3K7, Canada.

## Abstract

Fingertip force scaling during hand-object interactions typically relies on visual information about the object and sensorimotor memories from previous object interactions. Here, we investigated whether contextual information, that is not explicitly linked to the intrinsic object properties (e.g., size or weight) but is informative for motor control requirements, can mediate force scaling. For this, we relied on two separate behavioral tasks during which we applied transcranial magnetic stimulation (TMS) to probe corticospinal excitability (CSE), as a window onto the primary motor cortex role in controlling fingertip forces. In experiment 1, participants performed a force tracking task, where we manipulated available implicit and explicit visual information. That is, either the force target was fully visible, or only the force error was displayed as a deviation from a horizontal line. We found that participants’ performance was better when the former condition, in which they had explicit access to predictive information. However, we did not find differences in CSE modulation based on the type of visual information. On the other hand, CSE was modulated by the change in muscle contraction, i.e., contraction vs. relaxation and fast vs. slow changes. In sum, these findings indicate that CSE only reflects the ongoing motor command. In experiment 2, other participants performed a sequential object lifting task of visually identical objects that were differently weighted, in a seemingly random order. Within this task, we hid short series of incrementally increasing object weights. This allowed us to investigate whether participants would scale their forces for specific object weights based on the previously lifted object (i.e., sensorimotor effect) or based on the implicit information about the hidden series of incrementally increasing weights (i.e., extrapolation beyond sensorimotor effects). Results showed that participants did not extrapolate fingertip forces based on the hidden series but scaled their forces solely on the previously lifted object. Unsurprisingly, CSE was not modulated differently when lifting series of random weights versus series of increasing weights. Altogether, these results in two different grasping tasks suggest that CSE encodes ongoing motor components but not sensorimotor cues that are hidden within contextual information.

**Highlights:**

- Explicit visual cues and sensorimotor experience are key for object grasping
- Investigating influence of implicit, contextual information on object grasping
- Explicit but not implicit cues improve motor performance during object grasping
- Explicit but not implicit cues modulate corticospinal excitability

## 1. Introduction

Skilled hand object interactions require predictive planning of fingertip forces adjusted to the properties of the object (Gordon et al., 1993) as, for instance, a fragile object may be damaged when exerting too much force. Conventionally, it has been argued that individuals estimate object weight based on two mechanisms. First, by relying on visual cues and well-established “priors”, such as the size-weight (Gordon et al., 1991) and density-weight relationships (Baugh et al., 2012), object weight can be estimated. Second, when visual cues are not indicative of object weight (e.g., a suitcase that might be empty or filled) then individuals rely on a “sensorimotor memory”. This sensorimotor memory stores short-term information about object weight based on previous interactions with the same or similar objects (Baugh et al., 2012; Fu et al., 2010; Gordon et al., 1993; Johansson & Westling, 1988; Vonne van Polanen & Davare, 2015).

Interestingly, Cashaback et al. (2017) demonstrated that predictive object lifting not only relies on visual cues or sensorimotor memories, but also incorporates probabilistic information. In their study, participants minimized the potential prediction error across many lifts instead of optimizing their fingertip forces for the mostly anticipated object. In addition, Mawase & Karniel (2010) argued that an individual’s sensorimotor system can use information the individual is not aware of: The authors showed that individuals predictively scale their fingertip forces to a non-existent heavier weight after lifting a series of increasing weights that were implicitly presented within larger sequences primarily consisting of random weights. Other studies on hand actions have supported the notion that motor control mechanisms can adapt their sensorimotor predictions towards environmental variability for minimizing the general prediction error across multiple trials (e.g., Hadjiosif & Smith, 2015; Körding et al., 2004; Landy et al., 2012; Zhang et al., 2015). Importantly, it has been argued that finetuning motor commands to environmental variability relies both on explicit (i.e., conscious strategies) and implicit (i.e., unaware/subconscious adaptation) learning mechanisms (Albert et al., 2022). Notably, studies teasing apart explicit and implicit components of sensorimotor learning typically rely on reaching, and not grasping (e.g., Wolpert & Flanagan, 2016).

While it seems likely that sensorimotor learning during grasping relies on similar mechanisms as reaching due to the interconnectedness of their respective neural pathways (Gerbella et al., 2017), performance errors are typically perceived differently between reaching and grasping. During reaching, performance errors can be perceived by a visual mismatch between the target position and the actual end-position. During grasping, lifting errors are only perceived through a haptic mismatch between the expected and the actual object weight (Castiello, 2005; Johansson & Westling, 1984).

Due to (i) these differences between grasping and reaching, (ii) most of the scientific literature relying on reaching rather than grasping to understand implicit learning (for instance see Albert et al., 2022; Heald et al., 2021) and (iii) grasping studies typically relying on explicit visual cues to indicate object weight (e.g., Buckingham et al., 2014; Loh et al., 2010; Rens et al., 2021a), it is still unclear how the brain encodes implicit information for grasping. While it is plausible that the cortical grasping network (Davare et al., 2011; Gerbella et al., 2017) primarily relies on explicit visual cues and sensorimotor memories, it could still benefit from implicit learning. For instance, when lifting many opaque objects with varying weights (e.g., baggage handlers) it seems preferable to minimize the average prediction error than to try to predict each object’s weight (as in the study of Cashaback et al., 2017).

Importantly, the primary motor cortex (M1) plays a pivotal role in controlling hand-object interactions. When disrupting M1 activity transiently with repetitive Transcranial Magnetic Stimulation (rTMS), individuals fail to predictively scale their fingertip forces based on sensory feedback (Chouinard et al., 2005; Parikh et al., 2020; Schabrun et al., 2008). In addition, Loh et al. (2010) showed that corticospinal excitability (CSE), probed with TMS over M1 during lift planning, reflects the weight of the previously lifted object but becomes reflective of a forthcoming weight when indicated by a visual cue. In previous work from our group, we showed that CSE modulation can reflect an object’s weight distribution when indicated through action observation (Rens et al., 2021b). Arguably, these studies (Loh et al., 2010; Rens et al., 2021b) and similar ones (e.g., Baugh et al., 2012; Fu et al., 2010; for a review see Johansson & Flanagan, 2009) investigated how task requirements, indicated by explicit visual cues or sensorimotor memories are represented in corticospinal outputs during hand-object interactions.

To our knowledge, it is not well understood how CSE encodes hand-object interactions when task requirements are only implicitly present. For instance, in the study of Mawase & Karniel (2010), the question arises whether CSE modulation would represent force predictions based on an implicit series of increasing weights or the last lifted weight as in study of Loh et al. (2010). Noteworthy, using different behavioral modalities, CSE has been shown to be modulated by implicit (probabilistic) information. For instance, using a reaction time task, Van Elswijk et al. (2007) showed that CSE modulation increases with increasing response signal expectancy. That is, the longer it takes for the cue to appear, the more likely it is to appear, which increases CSE modulation. This finding was replicated by Dupin et al. (2019). Using a similar reaction time paradigm, Bestmann et al. (2008) showed that CSE modulation increases in case of low entropy (i.e., low uncertainty about sampling one trial from all possible trials) and low surprise (i.e., probability of the upcoming trial on itself). In another study, Senot et al. (2011) argued that, during lift observation, CSE is modulated by both written labels (i.e., explicit cues) and movement kinematics (i.e., implicit cues) indicating object weight. Similar results were found in another action observation study by Puglisi et al. (2017): CSE was modulated by explicit visual cues but also by implicit visual cues, i.e., in the authors’ words: “visual cues that were not the focus of attention”.

In the present study, we aimed to extend on studies investigating how environmental uncertainty affects grasping (see Cashaback et al., 2017; Hadjiosif & Smith, 2015; Körding et al., 2004; Landy et al., 2012) and explore how the sensorimotor system can integrate implicit information for grasping. For this, we used two separate experiments to investigate the integration of implicit information during continuous force exertion by squeezing (experiment 1) and during discrete object lifting (experiment 2). In addition to these behavioral tasks, we included TMS to probe CSE and investigate the underlying motor cortical mechanisms.

For experiment 1 we used a force tracking task and manipulated the type of visual information participants received about their ongoing motor performance. Participants were instructed to track a sinusoidal force curve by exerting increasing and decreasing force at different rates of change (slope of the curve) with their fingertips on force sensors. The type of the visual information was varied by displaying either the actual sinusoidal curve or a horizontal line. Importantly, in the sinusoidal curve condition, participants could rely on both predictive mechanisms, by observing the curve’s slope, and feedback-driven mechanisms, by observing their performance error (i.e., distance between the target curve and the line representing their performance). If participants would perform the task perfectly, their performance would overlap with the force target (sinusoidal curve). If they would not generate force at all, their performance would show as a horizontal line. In this way, we argued that both prediction- and feedback-related visual cues would be explicitly available. In contrast, in the horizontal line condition, participants tracked the same sinusoidal curve although the actual force target was hidden from the participants. Instead, they could only see their performance error (i.e., deviation from the sinusoidal curve) as deviations from a horizontal line. Thus, in this condition, if participants would perform the sinusoidal force tracking task perfectly, their performance would show as a horizontal line. Conversely, if participants would not exert force at all, their performance would be represented as the inverse of the sinusoidal target curve. As such, we argued that in this horizontal line condition, participants only had explicit access to their performance errors: they could only implicitly deduct their location on the hidden sinusoidal curve as (a) the force scaling requirements (curve slope) was not visible and (b) the performance error was shown in an inverse manner. Finally, TMS was applied to probe CSE during different phases (speed: fast or slow rate of change; target line: sinusoidal or horizontal; direction: up- or downscaling of fingertip forces). We argued that by manipulating the type of visual information, we could investigate how explicit and implicit information affects CSE modulation during visuomotor control of continuous hand-object interactions.

Using a similar paradigm, Kimura et al. (2003) asked participants to contract and relax their elbow in a sinusoidal manner. They found that CSE increased during the contraction compared to relaxation phase, even though TMS was applied when the same amount of force was applied in both phases. In line with Kimura et al. (2003), we hypothesised that, in both conditions, CSE would be larger during increasing compared to decreasing force exertion. To extend on the work of Kimura et al. (2003), we also manipulated the slope of the sinusoidal curve, resulting in slower or faster scaling of fingertip forces, to investigate sensorimotor involvement in the rate of force changes. We hypothesized that CSE modulation would be larger in the fast compared to slow condition. Importantly, if CSE modulation reflects ongoing task performance, independent of the type of visual information, CSE would be modulated similarly in the sinusoidal curve and horizontal line conditions. However, Bestmann et al. (2008) showed that CSE modulation is larger during low uncertainty. Accordingly, we hypothesized that, due to differences in the type of visual information between the target line conditions, CSE modulation would be larger when contextual information was more explicitly present, i.e., in the sinusoidal curve condition.

For experiment 2 we used an object lifting task that was similar to Mawase & Karniel (2010). Participants lifted small objects that were visually identical but had different weights. The total series of all lifted objects consisted of intermixed series of randomly ordered weights and series of increasing weights. Briefly, Mawase & Karniel (2010) showed that individuals predictively scale their fingertip forces to a non-existent heavier weight after lifting a series of increasing weights when this series was hidden within larger series consisting of random weights. Notably, in their study participants indicated during debriefing that they were unaware of these hidden series of increasing weights. As such, the authors argued that participants implicitly embodied the series of increasing weights, leading to extrapolation during predictive force planning. In this experiment, we argued that sensorimotor control mechanisms could extrapolate force requirements after the series of increasing weights, which would be reflected in changes in corticospinal output. That is, motor performance and CSE would not reflect a direct encoding of force scaling based on the last trials (i.e., the sensorimotor memory) but rather would reflect an extrapolation process for predicting force scaling based on (the implicit information) related to the series of increasing weight. As such, we hypothesized that participants would use more force when lifting the same weight after the implicit series compared to after the random series. Similarly, we hypothesized that CSE would reflect the extrapolated weight based on the series of increasing weights and not the weight of the previously lifted object (akin to Loh et al., 2010).

## 2. General Methods

### 2.1 Participants

Thirty healthy right-handed volunteers were recruited for the experiment, of which 15 took part in experiment 1 (12 females, mean age 22±2 years) and 15 in experiment 2 (11 females, mean age 22±3 years). Two participants (one for each experiment) were excluded from analyses, causing us to have 14 participants for each experiment. The first one (female) was excluded during data processing as the motor evoked potentials (MEPs; see below) during the experiment (and thus after defining stimulation intensity) exceeded the range of our electromyography (EMG) system. The second participant (female) was excluded as even during baseline assessment of CSE she was not able to relax their hand.

All participants were right-handed (self-reported). TMS eligibility was assessed by the TMS safety screening questionnaire (Rossi et al., 2011). Participants had no known neurological or sensory impairments and had normal or corrected-to-normal vision. The study was approved by the ethical committee of UZ/KU Leuven and all participants signed an informed consent before the experiment.

### 2.2 Acquisition of force data

Forces were measured using a grip(-lift) manipulandum (see Figure 1). The manipulandum consisted of two force/torque sensors (ATI Nano17, ATI Industrial Automation) that were mounted on a custom-made carbon fiber basket. The total weight of the manipulandum was 1.2 N. The graspable surface (17 mm diameter and 45 mm apart) of the force sensors was covered with fine sandpaper (P600) to increase friction. In the manipulandum’s basket, small objects could be placed. Force sensor data was sampled at 1 kHz frequency through a NI-USB 6343X (National Instruments, USA) which was connected to a personal computer. Measurement of forces in both experiments was performed using custom written Labview scripts (National Instruments, USA) and transferred for off-line analysis.

**Figure 1.**
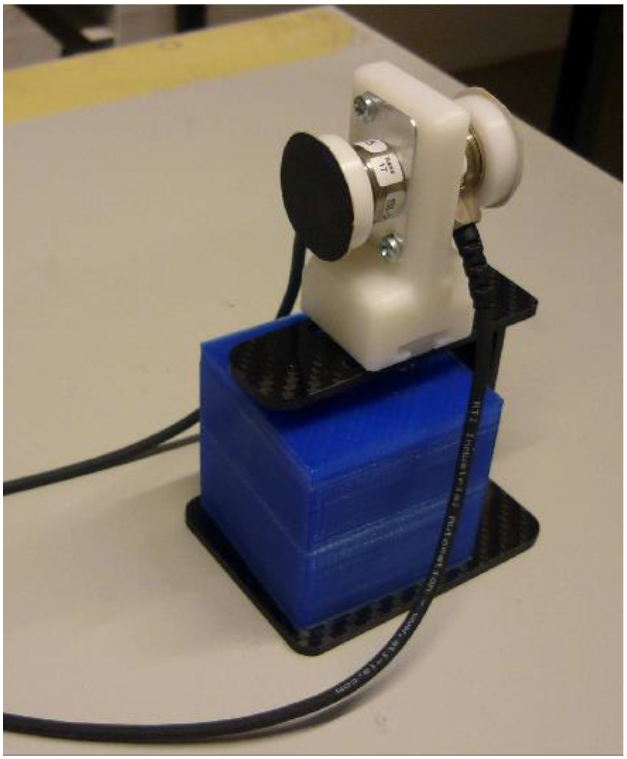
The (grip-lift) manipulandum consisting of force sensors mounted on a carbon fibre basket as used in the experiments. In experiment 1, participants squeezed the force sensors to track a curve on a computer screen. In experiment 2, small visually identical cubes with different weights were placed in the basket to be lifted by the participants.

### 2.3 Transcranial magnetic stimulation procedures

For both experiments, two Ag-Al electrodes were placed on the first dorsal interosseus muscle (FDI) in a typical belly-tendon montage. A third reference electrode was placed on the ulnar styloid processus. Electrodes were connected to a NL824 AC preamplifier (Digitimer) and a NL820A isolation amplifier (Digitimer), which in turn was connected to a micro140-3 CED (Cambridge Electronic Design). The CED was connected to a personal computer. Electromyography (EMG) recordings were amplified with a gain of 1000, high pass filtered with a frequency of 3 Hz, sampled using Signal and Spike2 software (Cambridge Electronic Design, UK) and were stored for off-line analysis. Spike2 was used for data collection in experiment 1 and Signal was used for data collection in experiment 2. The sample frequency was 1 kHz and 5 kHz in Experiment 1 and 2, respectively. A 70 mm figure-of-eight TMS coil [experiment 1: Magstim rapid 200 (Magstim Co Ltd, United Kingdom); experiment 2: DuoMag XT-100 (Deymed Diagnostic, Czech Republic)] was used to apply TMS. In both experiments, participants received TMS over M1 only. For M1 stimulation, the coil was tangentially placed over the head to induce a posterior-anterior current flow and to elicit motor evoked potentials (MEPs) in the FDI muscle. Before the experiment, we defined the optimal stimulation site (i.e., ‘hotspot’) and the stimulation intensity to be used in the experimental task for each participant. The hotspot was defined as the position from which MEPs with maximal amplitude were produced in the FDI muscle. The hotspot was marked on the scalp. The stimulation intensity (1mV threshold) was defined as the lowest stimulation intensity that produced MEPs equal to or greater than 1 mV in the FDI muscle in at least five out of ten consecutive trials when stimulating at the predetermined hotspot.

#### Specific procedures for experiment 1

The behavioural task of experiment 1 consisted of continuous force production (i.e., ‘squeezing’ on the F/T sensors; see below). We assessed the 1 mV threshold while participants produced a constant total grip force of 2.5 N (sum of horizontal forces applied by thumb and index finger on the force sensors). As such, the 1 mV threshold was not defined during ‘rest’ but during active contraction. The 1 mV threshold for experiment 1 was 42 ± 11 % (group mean ± standard error) of the maximal stimulator output.

#### Specific procedures for experiment 2

The behavioural task of experiment 2 consisted of an object-lifting task and TMS was applied during lift planning (i.e., before the reaching phase, thus while the hand was still relaxed on the table; see below). Accordingly, the stimulation intensity for experiment 2 was defined while participants kept their hand relaxed on the table. The 1 mV threshold was 56 ± 8 % (group mean ± standard error) of the maximum stimulator output.

## 3. Methods of Experiment 1

### 3.1. Task and procedures

After initial preparations (i.e., consent, questionnaires and TMS preparations), participants performed the experimental task. Participants were comfortably seated in a chair. A table was positioned in front of them on which a computer screen was placed and the grasp-lift manipulandum was placed on another table on their right side. Participants were instructed to rest their right arm on the table and grasp the two force sensors of the manipulandum using their right thumb and index finger (precision grip). The manipulandum itself was fixed on the table to ensure it would not move during the experiment. During the experiment, the TMS-coil was held above the hotspot by an experimenter (see Figure 2 for a drawing of the set-up).

**Figure 2.**
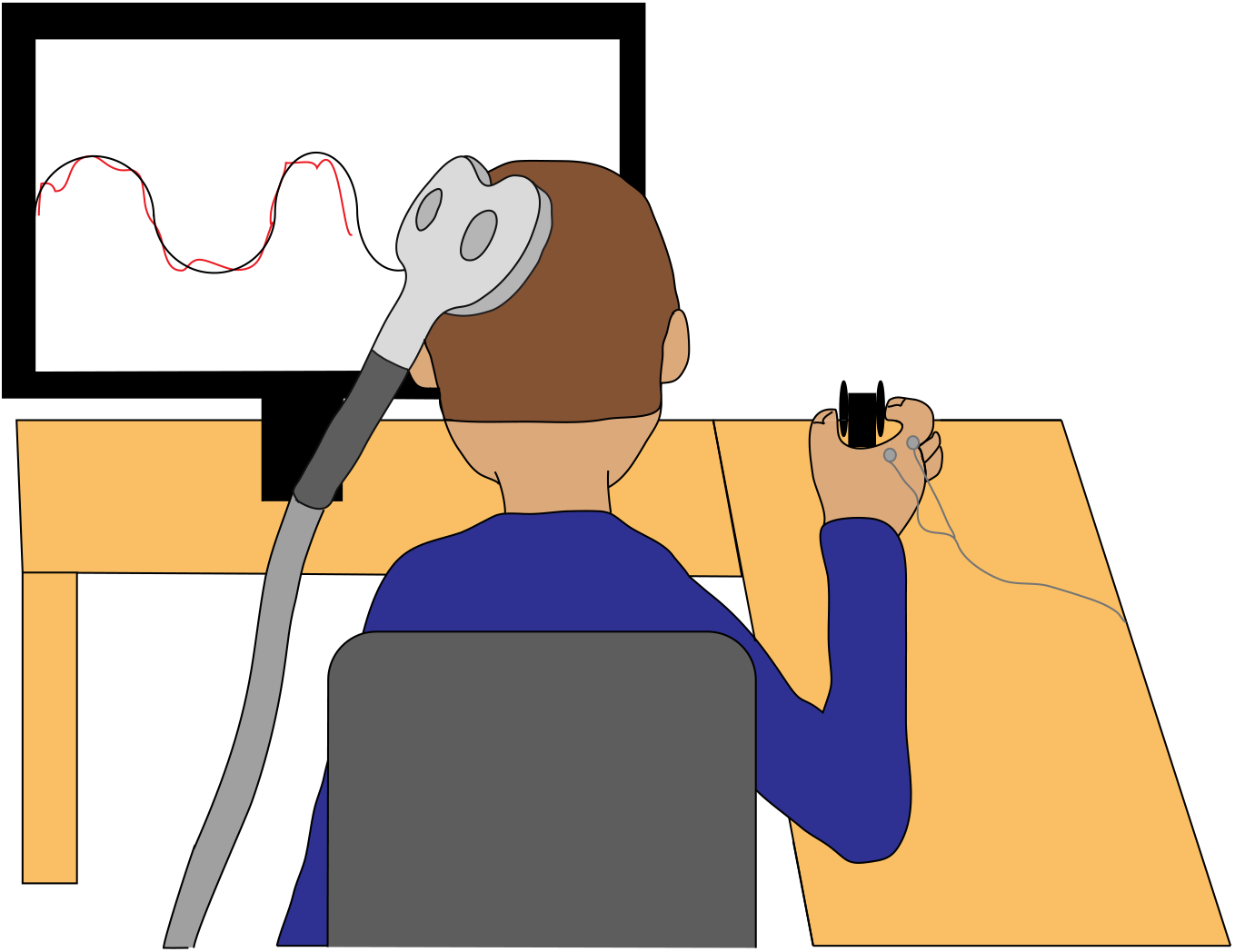
Drawing of the experimental setup in experiment 1. Participants squeezed the force sensors to track a target curve on the screen with their right hand. During the task, transcranial magnetic stimulation was applied over the left motor cortex and electromyography was measured with electrodes placed on the right first dorsal interosseus muscle.

#### Sinusoidal curve condition

On the screen a sinusoidal target line was projected with the x- and y-axis representing time and amount of exerted force, respectively (Figure 3B). The target sinusoid had an amplitude of 1.2 N and was centred around 2 N. Given that we wanted to avoid muscle fatigue for the TMS data (Kotan et al., 2015), we decided to have a light muscle contraction (2 N average) and relatively small change in force exertion (1.2 N sinusoidal increase/decrease). The 2 N was arbitrarily defined close to the lightest weights used in our previous studies (e.g., 1.5 N in Rens et al., 2020, and 2.2 N in Van Polanen et al. 2020). The rationale was that we found clear MEPs when participants planned to lift these weights in those studies. As such, we argued that we could investigate a low-intensity muscle contraction during squeezing (this study) of which we could still anticipate MEP effects based on our previous object-lifting work. The change in force exertion (1.2 N sinusoidal increase/decrease) was arbitrarily defined as 60 %of 2 N. Given that we applied TMS only above 2 N, task-related MEPs should have been present. A cursor moved over the target line indicating the participant’s location over time. The cursor produced a red line providing visual feedback of performance over time. Participants controlled the cursor’s vertical displacement by exerting force on the force sensors. Participants could not control the cursor’s horizontal displacement as the cursor moved with a constant speed over the screen. The sinusoid could have a frequency of either 0.09 (‘slow’ condition) or 0.25 Hz (‘fast’ condition) over the positive and negative half cycles which was indicated by a ‘wider’ or ‘smaller’ sinusoidal wave thus causing a more gradual or steep slope, respectively. The frequency (thus the sinusoid’s width) was changed pseudo-randomly when crossing the midpoint (i.e., when exerting a force equal to 2 N). In this way, four different conditions could be identified, being fast up (upscaling of forces; 0.25 Hz), fast down (downscaling; 0.25 Hz), slow up (upscaling; 0.09 Hz) and slow down (downscaling; 0.09 Hz). Our frequencies were based on Kimura et al. (2003) who used three frequency conditions (slow: 0.08 Hz, medium: 0.17; fast: 0.33Hz). Because we included different visual conditions (sinusoidal curve vs horizontal line) as well, we decided on two frequency conditions instead of three. Based on pilot experiments to verify comfortable speeds for our squeezing task, we chose frequencies of 0.09 and 0.25 Hz. Note that we did not intend to directly compare our results with Kimura et al. (2003) as we were not particularly interested in the relationship between MEP amplitude and contraction speed. Instead, we were mainly interested in relative differences (i.e., how do MEPs differ between faster and slower contractions) as basis to explore the effects of different types of visual information. Participants were instructed to stay as close as possible to the target line by controlling the vertical displacement of the cursor and, thus, by sinusoidally changing the amount of force they were exerting on the force sensors. As participants could see where they were on the target line, they could anticipatorily plan to up- or downscale their forces and their speed. Optimal performance in this task would consist of sinusoidally scaling fingertip forces and would show the red trace (produced by the cursor’s movement) to perfectly overlap with the sinusoidal target line. Last, TMS was applied when participants exerted 2.5 N (Figure 3B, green asterisk). As such, TMS was only applied during the positive half-cycle (considering that the sinusoid was centered around 2 N). We argued that we could exclude stimulating during the negative half cycle. Inclusion of these timings would only allow us to additionally compare the amounts of exerted force which did not add to our research questions. In addition, excluding TMS during the negative half cycle allowed us to reduce experimental duration and to minimize potential confounding effects of muscle fatigue and/or decline in attention. We did decide to stimulate during both upward (increasing contraction) and downward force scaling (increasing relaxation) during the positive half cycle to validate our findings with respect to Kimura et al., (2003). In addition, we arbitrarily decided to stimulate at 2.5 N based on the following rationale: in the fast condition (0.25 Hz), a cycle is completed in 4 seconds. As such, from the midpoint to the first peak takes approximately 1 second (one fourth of cycle). If TMS is applied after a 0.5 N increase (± 40 % of maximum increase of 1.2 N), then TMS should be applied earliest ± 400 ms after reaching the 2 N midpoint in the fast condition. Based on grasping studies (for a review see: Castiello, 2005), we argued that this would enable participants to integrate feedback if their performance “faltered” after switching conditions (in particular from slow to fast). To end, the interval between two TMS pulses was minimally 3.5 seconds. The latency between TMS pulses should not have resulted in carry-over effects on MEPs (e.g., longer-term inhibition) (Cincotta et al., 2003; Hannah, 2020; Wilson et al., 1993).

**Figure 3.**
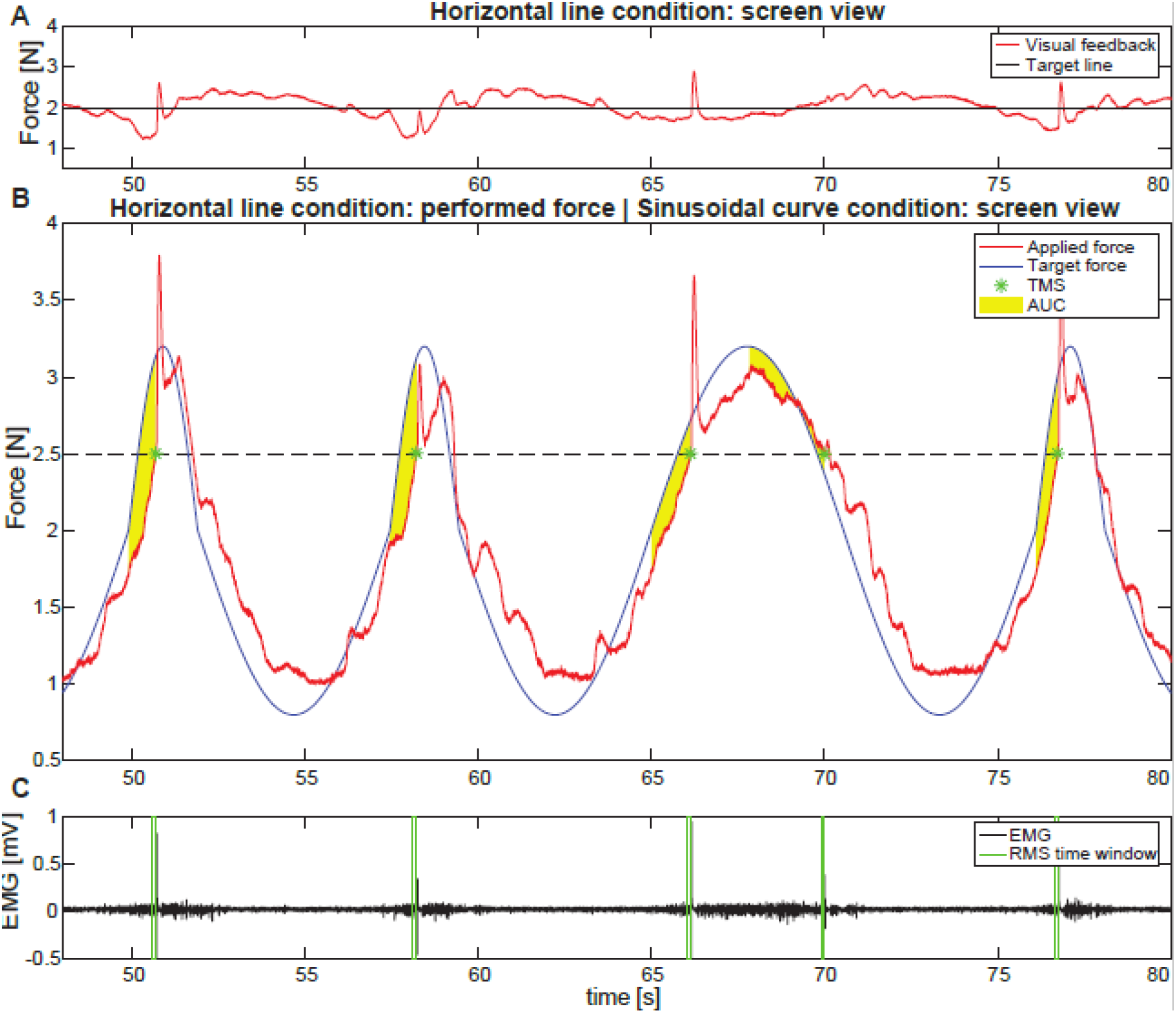
Representative example of a part of a trial in the horizontal line condition. **A.** Screen view as shown to the participant during performance of the task in the horizontal line condition. The black line indicates the target line and the red line represents the participant’s performance as deviation in exerted force from the target line (in N). **B.** Actual performed force (red) and actual target force which was hidden from the participant (blue). The black dashed line represents the force level at which TMS was applied. A TMS event is indicated with a green asterisk. Yellow shadings indicate the area under the curve (AUC) as a measure of performance. Note that in the sinusoidal condition, the screen view looked like the blue (target) and red (performance) lines. **C.** Electromyography (EMG) data (black). The time window of 100 ms before TMS that was used to calculate the background EMG (RMS100) is indicated with green vertical lines.

#### Horizontal line condition

The experimental design was nearly identical to the sinusoidal curve condition. As such, we will only discuss differences. In this task, the sinusoidal target line was not shown on the screen, but only a horizontal line was shown (Figure 3B). Participants had to perform the same task of sinusoidally scaling their force. The cursor showed the deviation from the ‘hidden’ sinusoidal target curve as a deviation from the horizontal line. Accordingly, participants could not perceive where they were (e.g., positive, or negative half cycle of sinusoidally scaling their forces), but they could still perceive the red trace produced by the cursor’s movement. As such, participants could not anticipatorily plan their forces (e.g., by seeing that they were at the peak of the positive half cycle or whether they were performing a cycle in the slow or fast condition) but they still received on-line feedback about their performance. Optimal performance in this task would consist of sinusoidally scaling fingertip forces but would show the red trace (produced by the cursor’s movement) overlapping with the straight horizontal target line.

The top row (Figure 3A) shows a representative example of a participant’s on-screen view and their performance in the horizontal line condition. The sinusoidal curve was hidden (i.e., only a target horizontal line was shown) although participants still received visual feedback about their ongoing performance. The second row of Figure 3 (Figure 3B) shows the actual sinusoidal curve, that was hidden from the participants, and the participant’s performance relative to this sinusoidal curve. When a participant would perform the sinusoidal curve condition, the target curve and the feedback of the participant’s performance would look like the example in Figure 3B. On this figure it can also be seen that in this example the first two and last cycles are part of the ‘fast’ condition, whereas the third cycle is part of the ‘slow’ condition.

**Figure 4.**
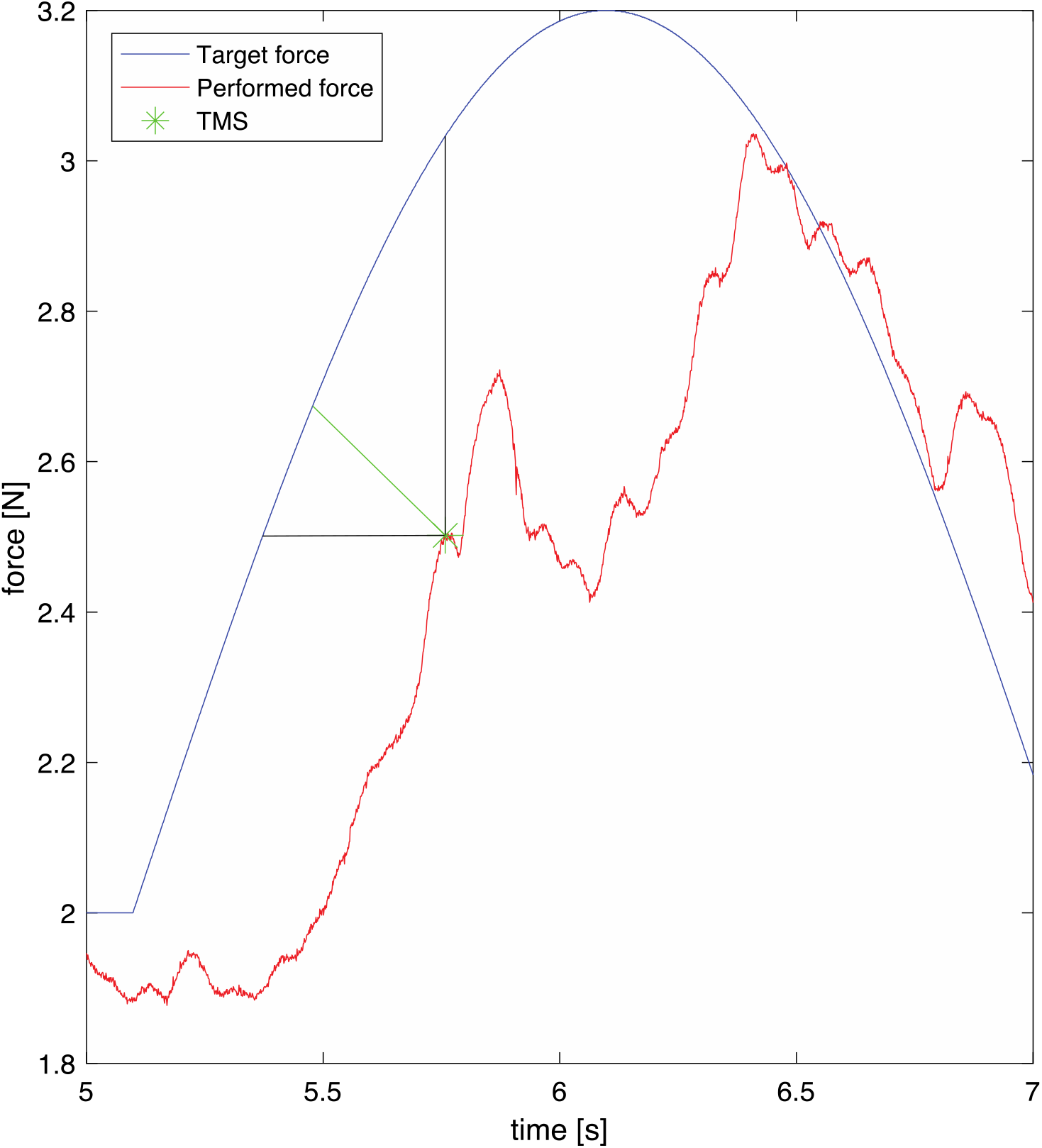
Example of calculation of Euclidian distance to determine erroneous trials. The blue and red line represent the target and performed force, respectively. From the point at which the TMS was applied (green asterisk) the shortest distance to the target force was determined, which was done in 2D space incorporating both the deviation in time and force. Black lines indicate distances to the target curve if only force or time were considered.

The sinusoidal curve and horizontal line conditions were presented in a blocked design which were counterbalanced across participants to avoid order effects. In each block, 13 trials of 30 seconds were performed (6 participants) or 5 trials of 120 seconds (8 participants). The length of trials was changed after the first six participants as it became apparent that participants could easily perform trials of longer durations. Each trial consisted of a random sequence of the four conditions (frequency: slow-fast; direction: down-up). In each trial, each condition was equally presented to avoid potential fatigue and decline-in-attention effects. To ensure that group identity (i.e., either performing short or long trials) did not impact task performance, we compared performance (AUC parameter as described in Data analysis) for each condition between our groups using student t-tests. For each comparison (e.g., slow up of the short trial group vs slow up of the long trial group), we found a p-value of minimally p = 0.144 which are not corrected for multiple comparisons. Because of this, we argued that there are no meaningful differences between both groups and, thus, decided to pool both groups for further analyses.

At the start of each trial, a preparatory target line at the 2 N level was shown. The cursor took between 5 to 8 seconds (randomized) to complete, allowing participants to move the cursor to the midpoint force level at the start of the trial. This preparatory target line would then transition into the experimental sinusoidal curve or horizontal line. The transition between the preparatory and experimental target line was indicated with a vertical line. Before starting the experimental task, 2 practice trials of 30 seconds were performed, and a short break was presented between the two blocks.

### 3.2 Data analysis

Several parameters were extracted when TMS was applied (i.e., when participants generated 2.5 N); the MEP amplitude (as defined above) was extracted in a window of 5 to 45 ms after the TMS pulse. From the EMG, we also extracted muscle activity using peak-to-peak amplitude and root mean squared activation (RMS100) in a 100 ms interval before the TMS pulse (105-5 ms before the TMS pulse) as a measure of background EMG before TMS.

Although participants were instructed to keep the cursor as close as possible to the target line, they often made errors. Particularly in the horizontal line condition, participants could not rely on visual information regarding the upcoming force requirements as they could only rely on their ongoing performance (as indicated by increasing/decreasing deviations from the target horizontal line). Accordingly, participants sometimes scaled their fingertip forces not in line with the current event (e.g., up-instead of downscaling or fast instead of slow scaling). To assess (i) which event should have been performed (up/down – fast/slow) and (ii) performance on this event, we did the following: we calculated the total error in 2D space (force-time space). This allowed us to include errors in force (too much/too little force applied at any given give) and time (where participants should have been on the curve when applying a certain amount of force). The total error for each event was defined as the shortest Euclidean distance between the target curve and the participant’s force generation at the TMS event (see Figure 4). Specifically, in Figure 4, the TMS event is indicated with a green asterisk on the actual performed force (red line). The shortest distance from this asterisk to the target curve (i.e., between current performance and the target sinusoidal curve in blue) was determined. Note that in the horizontal line condition, even though participants perceived a straight horizontal target curve, we used the ‘hidden’ sinusoidal curve to investigate performance (i.e., participants saw on-screen Figure 3A, but their performance was quantified as in Figure 3B). We decided to include both dimensions (force and time) for assessing general performance as, for instance, participants could have been generating the appropriate amount of force although they were performing the incorrect event (e.g., up-instead of downscaling forces). As the TMS pulse was automatically triggered when participants generated 2.5 N (with a minimum 3.5 s delay between two consecutive stimulations), we had no direct control over when TMS was applied. We reasoned that if participants deviated too much in force-time space, the MEP would not reflect the current event and thus needed to be removed from analysis. Based on visual inspection of all trials, a criterion of 0.55 was used for this total error. This value of 0.55 was chosen as it closely matched the visual judgement of two observers that independently rated the individual trials. Using this criterion, 2% of all MEPs were removed. Note that we did not include this parameter (the Euclidian total error) to assess participants’ overall performance but rather to reject incorrectly performed events. In addition, we also removed MEPs that were smaller than the peak-to-peak EMG in the 100 ms interval before TMS (134 trials, 4%), as these MEPs lower than background EMG activity likely reflect noise.

Last, MEPs were obtained from TMS applied at the same force level. Since EMG amplitude depends on the amount of force that is applied, the RMS100 should not differ between the conditions. However, we found that this was not always the case (see Results). To control for differential force activation, we also divided the MEP values by the RMS100 before the TMS pulse and analysed this corrected value (MEP_EMG_) separately. We decided to perform this normalization to be in line with Kimura et al. (2003) and given that MEP amplitude varies with muscle contraction (Lavoie et al., 1995).

To investigate actual participants’ performance, we relied on force-related parameters. The average force rate (time differentiated force) was determined between 300 ms before the TMS pulse and the TMS pulse. Furthermore, to examine the deviation from the target line for a longer period, the difference between the target force and the generated force was calculated from the midpoint of the curve (start of a condition) until the time point TMS was applied. The absolute area under the curve of this difference (AUC) was calculated as a measure of deviation from the target curve. To correct for possible different time durations the AUC was divided by the number of samples included in its calculation. In this way, the AUC represented the average deviation between the executed force and the target force from the start of the curve to the time of TMS in a specific condition. Finally, to further evaluate task performance, the average speed was calculated as the mean force rate in the 300 ms before the TMS pulse.

### 3.3 Statistical analysis

All variables (MEP, RMS100, average speed, AUC) were calculated for each condition (i.e., fast up, fast down, slow up or slow down) and each task (sinusoidal curve or horizontal line). Since TMS was applied 3.5 s apart and always at the performed force level, the number of events in each condition differed among participants (as shown in Table 1 and explained in calculating the total Euclidian error in ‘Data Analysis’). A repeated measures analysis of variance (ANOVA) was performed on the variables with factors direction (up, down), speed (slow, fast) and task (sinusoidal curve or horizontal line). In line with previous work of our group (Rens et al., 2020), we z-scored MEP and RMS100 data for each participant separately before calculating participant averages and performing statistics. Post-hoc tests were performed with Bonferroni corrected t-tests. A significance level of 0.05 was used and effect sizes for the factors are reported using partial eta squared (*η*_p_^2^). Only if otherwise denoted, Data in text is reported as mean ± standard error.

**Table 1.** Events included in the analysis for each condition on average (after removal of incorrect events). Mean ± SD over participants for each condition.

|  | Slow up | Fast up | Slow down | Fast down |
| --- | --- | --- | --- | --- |
| Sinusoidal curve | 25.3±4.8 | 28.9±5.6 | 24.0±7.2 | 18.7±6.6 |
| Horizontal line | 26.1±6.0 | 26.2±8.3 | 21.7±8.6 | 17.5±5.0 |

## 4. Methods of Experiment 2

### 4.1 Task and procedure

After initial preparations (i.e., consent, questionnaires and TMS preparations), participants performed the experimental task. The behavioural task of experiment 2 comprised of an object-lifting task based on Mawase & Karniel (2010). Participants were comfortably seated in front of a table with their right arm resting on the tabletop. The experimenter was seated in front of the participant. Participants were instructed to grasp and lift the manipulandum on the table and received the following instructions: (1) Lift the manipulandum to a height of approximately 5 cm at a smooth pace that is natural to you. (2) Only place thumb and index finger on the graspable surfaces (i.e., use precision grip). (3) After lifting, keep the manipulandum in the air until the switchable screen (see below) turns opaque. Only then replace the object on the table. (4) Only start reaching towards the manipulandum after you have received the TMS pulse. Accordingly, one trial consisted of one reach-grasp-lift-replace action. Before the start of the experiment, five practice trials were performed with a random sequence of objects included in the experimental task.

A switchable screen (MagicGlass) was placed between the participant and the table. The screen could become transparent or opaque to enable/block vision on the manipulandum, respectively. When the screen was opaque between trials, the experimenter could change the cube in the manipulandum’s basket out of the participant’s view. Five possible objects could be placed in the manipulandum with approximate weights of 1.04, 2.55, 3.11, 3.56 and 5.15 N, which will be referred to as (weight) 1-5. As mentioned before, the manipulandum weighted approximately 1.2 N, thus giving total weights of approximately 2.23, 3.75, 4.31, 4.75 and 6.34 N to be lifted. Total weight was assessed by holding the cube-loaded manipulandum steady in the air for a few seconds which was repeated 5 times for each weight. These cubes were 3D printed, visually identical and measured 5×5×5 cm. In addition, all objects were hidden below the same paper cover to ensure that participants could not rely on visual cues indicating object weight.

Each trial started with the switchable screen becoming transparent. After 400 ms (± 100 ms jitter), a TMS pulse was applied over M1. Four seconds after the screen becoming transparent, it turned opaque again indicating the end of the trial and indicating participants to replace the manipulandum on the table. Force data was only sampled when the screen was transparent, EMG data was continuously sampled but only an epoch of 5 seconds (from 2 seconds before to 3 seconds after the TMS pulse) was stored.

The TMS pulse served as the ‘go’ cue for participants to initiate their action of grasping, lifting, and replacing the manipulandum. We decided to stimulate during planning, not execution, as we did not want to interfere with the participants’ lifting performance which would be caused by the unvoluntary muscle contraction due to M1 stimulation. To ensure that participants could not anticipate the TMS timing, we decided to include a jitter of 100 ms. We decided to use 300 ms as the lower threshold (thus 400 ms ± 100 ms jitter) as this would provide minimally 300 ms of visual information to participants. Although TMS during lift planning was applied substantially later in our study than in the one of Loh et al. (2010), it should still elicit CSE modulation as shown in previous work of our group (Rens et al., 2021b). In addition, Parikh et al. (2014) used a similar object lifting paradigm, applying TMS 1000 ms after object presentation. Briefly, in their study, CSE was also task-specifically modulated. Altogether, these studies indicate that TMS at the ‘go cue’ probes planning effects on CSE modulation, irrespective of the timing of the cue itself.

Similar to the task of Mawase & Karniel (2010) the weights were presented in an apparently random order although we used both series of increasing weights and of random weights. A series of increasing weights consisted of the weights 1, 2, 4 and 5 (i.e., four trials). After this series of increasing weights, weight 3 was presented in the fifth trial as a ‘catch’ trial’. The series of random weights varied between 4 and 9 trials of which a randomly chosen trial comprised of lifting weight 3 (weight 3 was presented earliest trial 2 and latest trial 9). This was to ensure that participants would not anticipate that the catch trial was the fifth trial after the series of increasing weights.

As weight 3 was presented after both the series of increasing and of random weights, we could compare force scaling in catch trials between these two conditions. Accordingly, we could investigate whether participants would predictively scale their forces differently for the same weight based on implicit information (i.e., increasing vs. random series). Importantly, to ensure that predictive force scaling differences for weight 3 between series were not driven solely by the previous lift (Johansson & Westling, 1988), we decided that the weight immediately before weight 3 in the series of random weights was the same as the last weight in the series of increasing weights (i.e., weight 5) (for an example sequence of the object lifting task see Figure 5). For experiment 2, we decided on 15 catch trials per condition, thus resulting in 15 series of increasing weights and 15 series of random weights which was close to double the amount of trials in Mawase & Karniel (2010).

**Figure 5.**
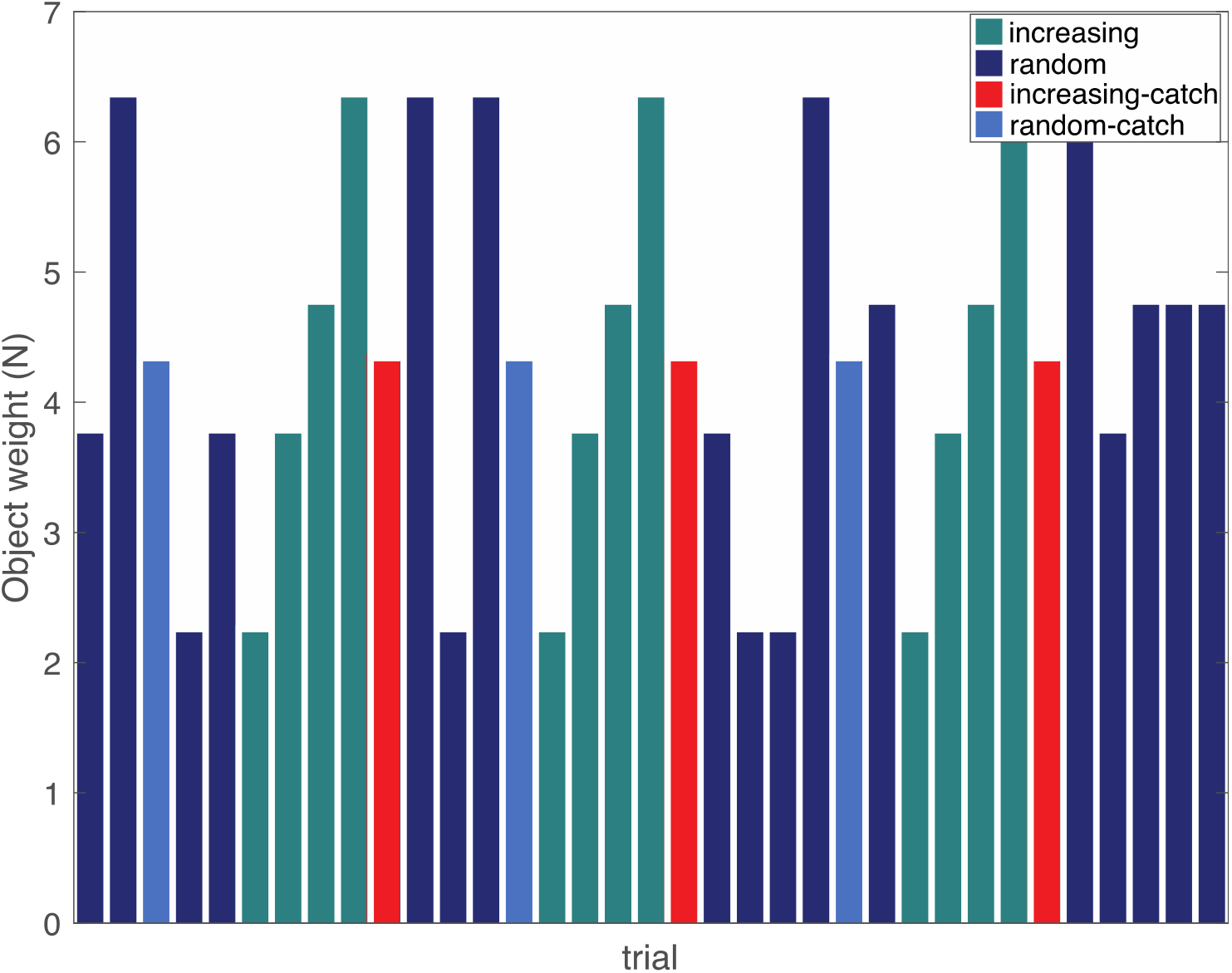
Example of trial sequence in experiment 2. Trials consisted of a series of increasing weights (turquoise) or random order (dark blue). A catch trial was presented in each series: at the end of an increasing series (red) or at a random position in the random series (light blue). Note that a catch trial was always preceded by the heaviest weight in the object set.

A different experimental sequence was generated for each participant. The 15 series of increasing weights included 4 weights and a catch trial, giving a total of 15 x 5 = 75 trials. The 15 random series consisted of 4-9 random weights (including a catch trial), giving on average 6.5 x 15 = 97.5 trials. In addition, each experimental sequence included 5-10 and 2-4 random trials at the start and end of the experiment, respectively. In total, participants performed on average 181 trials. Participants received a break after approximately 60 and 120 trials which varied as the break took place after completing a series (thus after lifting weight 3). After completing the experiment, participants performed a short questionnaire (akin to Mawase & Karniel, 2010), which contained questions about their experience of the objects:

1. Did you experience a change in the weight of the objects?
2. Did you experience this change in object weights as random?
3. Did you experience a pattern or series in the change of object weights?
4. Did you feel like some weights appeared more often or less frequent, or was this equally divided?

### 4.2 Data analysis

#### EMG data

From the EMG recordings, we extracted the MEPs, which were defined as the peak-to-peak amplitude in the EMG signal, using a custom-written MATLAB script. MEPs were determined between 5 ms to 45 ms after the TMS pulse. Trials were visually inspected for background noise related to muscle contractions. Trials that showed visual contamination of EMG activity prior to TMS were flagged and removed from analysis (3.6 %). In addition, MEPs were also considered noise and removed when they were smaller than 50 μV or smaller than the peak-to-peak EMG signal in the 100 ms before TMS (1.7 %) as they could not be discerned from background noise.

#### Force data

Trials in which the incorrect object was lifted (i.e., experimenter placing the wrong weight), or in which the object was not lifted or lifted twice in one trial (e.g., when the participant’s fingertips slipped from the force sensors) were removed from the force analysis (0.3%). The force data was filtered with a fourth order low-pass bidirectional Butterworth filter with a cut-off frequency of 15 Hz. This filter was selected to provide an optimal balance between noise removal and signal smoothing to find the peaks in the force rates (Van Polanen et al., 2020; van Polanen & Davare, 2015). Total load force (LF) and total grip force (GF) were the sum of the forces parallel and perpendicular to the gravitational force, respectively. Load and grip force rates (LFR and GFR) were defined as the first time-derivatives of LF and GF. Peak values of the force rates were determined between object contact (defined as GF > 0.1 N) and 50 ms after lift-off (LF > object weight). We decided to use the peak values, and in particular the first peak values (peak1 LFR and peak1 GFR), of the force rates (and not the forces) as this has been considered a more sensitive measure of predictive force scaling (Johansson & Flanagan, 2009; Johansson & Westling, 1988). In line with Van Polanen et al. (2020), we extracted the first peak that was at least 30% of the maximum value to avoid noise extraction. Finally, the loading phase duration (LPD) was determined as the latency between LF onset (LF > 0.1 N) and object lift-off (defined above). To end, we also calculated grip force at lift-off (GFatLO). GFatLO was extracted as this was the behavioural parameter used in (Mawase & Karniel, 2010). This allowed us to directly compare our findings with theirs.

### 4.3 Statistical analysis

We extracted all parameters (MEP, GFatLO, peak 1 GFR, peak1 LFR, LPD,) for catch trials after the series of increasing and random weights. For each parameter, outliers, defined as being smaller or larger than the mean ± three standard deviations, were removed from the analysis (< 2 % for any parameter). Analyses were performed on the averaged participant data. In line with Rens et al. (2020), MEP data were converted to z-scores to take inter-subject variability in account. Data were then averaged per parameter for each participant and condition and then used for statistical analysis. While data was obtained for longer series (see Figure 7), we only performed contrast comparisons on the last (catch trial) and the second to last lift (weight 5) of the series of increasing and random weights. For this, we used a repeated measures ANOVA with two factors being series (increasing vs random) and weight (weight 5 vs catch trial). Note that we only included the last two lifts due to the series of increasing weights consisting of 5 lifts and the series of random weights of at least 3 and maximally 5 lifts. In addition, while the last two lifts of both series were always the same (weight 5 and catch trial), the second to last lift of the random series varied randomly, providing no meaningful inputs for the statistical analysis. Post-hoc tests were performed with Bonferroni corrected t-tests. A significance level of 0.05 was used and effect sizes for the factors are reported using partial eta squared (*η*p^2^). Only if otherwise denoted, data is reported as mean ± standard error. To end, we calculated correlations between the weights and the participant averages for each parameter (MEPs and behavioral). This was done to explore a potential linear relationship and was performed once including the catch weight and once excluding it.

**Figure 6.**
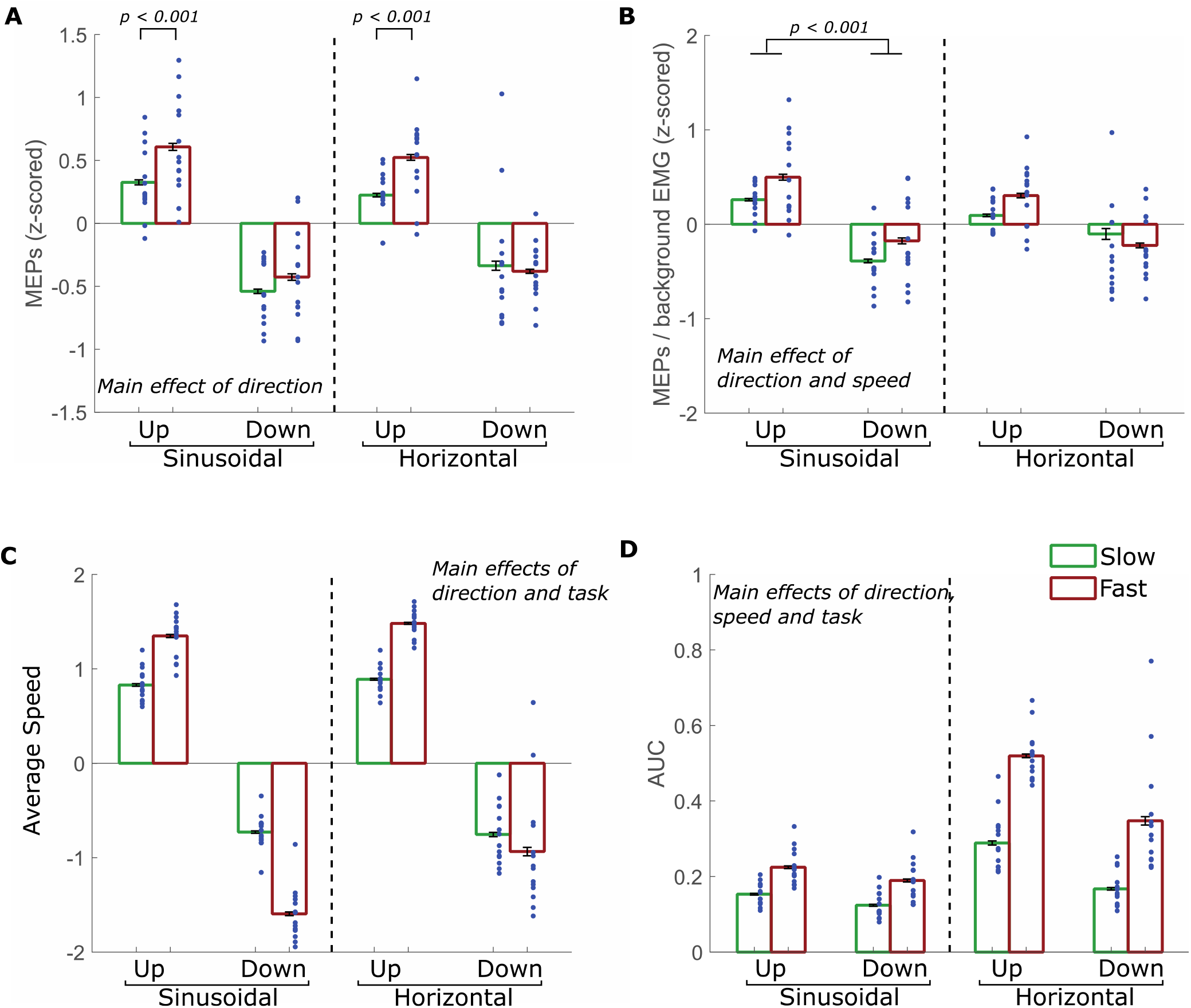
Results for experiment 1, for both the sinusoidal and horizontal line condition. **A.** Motor evoked potentials (MEP). **B.** MEPs divided by background EMG. **C.** Average speed. **D.** Area under the curve (AUC). Green and red bars represent slow and fast conditions, respectively. Blue dots represent individual values. Main effects are reported and interactions are indicated with significance bars. Error bars indicate standard errors.

**Figure 7.**
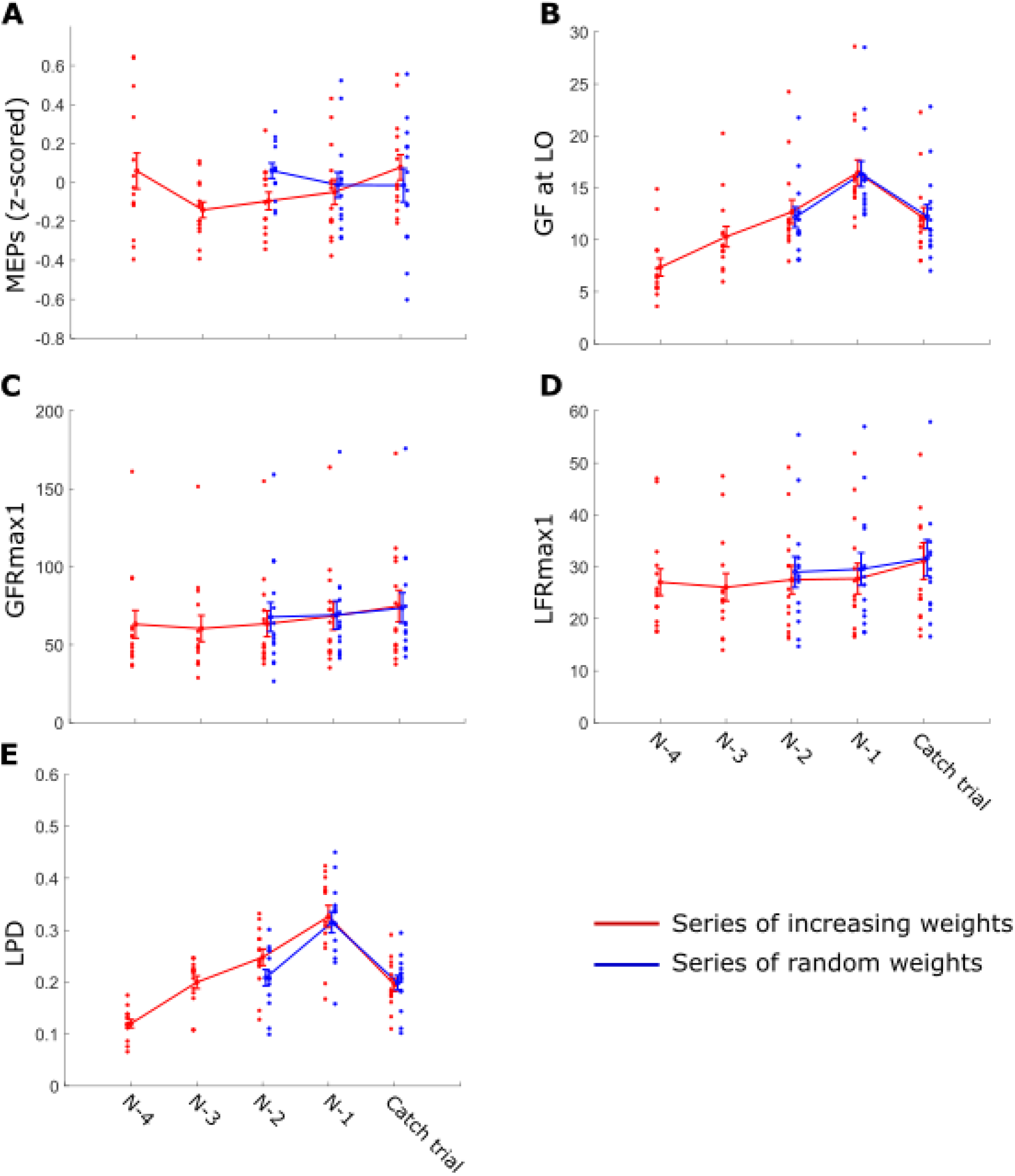
Results for experiment 2 for the catch trial after a series of increasing (red) or random (blue) weight. X-axis represents the catch trial (trial N) and all preceding trials for each series. For the series of increasing weights, the catch trial was preceded by four increasing weights (weight 1,2,4 and 5 which are N-4, N-3, N-2, and N-1 respectively). For the series of random weights, the catch trial was preceded by weight 5 (trial N-1) and minimally by at least one random weight (trial N-2). **A.** Motor evoked potentials (MEPs), z-scored. **B**. Grip force at lift off (GFatLO). **C**. first peak of grip force rate (GFRmax1). **D.** first peak of the load force rate (LFRmax1). **D**. Loading phase duration (LPD). Error bars indicate SEM. Colored scatters indicate the average of each participant for each condition.

## 5. Results

### 5.1 Experiment 1

In the first experiment, participants tracked a curve by applying force on two sensors. TMS was applied when a force of 2.5N was crossed, which could occur when moving up or down on the curve and at a fast or slow speed.

#### Corticospinal excitability

The MEPs were compared for all conditions to test whether CSE differed depending on the task that had to be performed. Results are shown in Figure 6A. The repeated measures ANOVA showed that MEPs were larger when moving up (up = 0.42 ± 0.04) compared to moving down (down = −0.42 ± 0.07) on the curve (effect direction, F(1,13) = 74.9, p<0.001, *η*_p_^2^ = 0.85). Furthermore, MEPs were also larger when the movement was fast (fast = 0.08 ± 0.03) compared to slow (slow = −0.08 ± 0.04) (effect speed, F(1,13) = 22.8, p<0.001, *η*_p_^2^ = 0.64). However, an interaction effect for speed x direction (F(1,13) = 7.8, p = 0.015, *η*_p_^2^ = 0.38) indicated that the speed effect was only seen in upwards movements (up fast = 0.57 ± 0.06; up slow = 0.27 ± 0.03; p = 0.002), but not in downwards movements (down fast = −0.40 ± 0.07; down slow = −0.44 ± 0.09; p = 1.00). The difference in CSE modulation was also seen between moving up and down within both the fast (up fast compared to down fast) and slow (up slow compared to down slow) movements (both p<0.001). Finally, an interaction for direction x task was found (F(1,13) = 7.2, p = 0.019, *η*_p_^2^ = 0.36), but post-hoc tests only showed differences between up and downward movements within tasks (sinusoidal curve: up = 0.47 ± 0.06; down = −0.48 ± 0.03; p < 0.001; horizontal line: up = 0.37 ± 0.07; down = −0.36 ± 0.1; p < 0.001), but no differences within direction between tasks (e.g., upward movement compared between the sinusoidal curve and horizontal line conditions) were found (both p = 1.00). To end, the main effect of task (F(1,13) = 0.35, p = 0.855, *η*_p_^2^ = 0.003) as well as the interaction effects of task x speed (F(1,1) = 0.43, p = 0.524, *η*_p_^2^ = 0.032), and task x direction x speed were not significant (F(1,13) = 0.70, p = 0.418, *η*_p_^2^ = 0.05).

#### Background EMG

Since MEPs also depend on background EMG, we compared the RMS of the EMG obtained 100ms before the TMS was applied (RMS100). Here, an effect of direction (F(1,13) = 112.90, p<0.001, *η*_p_^2^ = 0.90) was found. RMS100 was higher when going up (up = 0.28 ± 0.04) compared to going down (down = −0.44 ± 0.04). The main effects of task (F(1,13) = 0.53, p=0.481, *η*_p_^2^ = 0.039) and speed (F(1,13) = 2.37, p=0.148, *η*_p_^2^ = 0.15) were not significant as well as the interaction effects of task x direction (F(1,13) = 0.11, p=0.751, *η*_p_^2^ = 0.008), task x speed (F(1,13) = 0.62, p=0.45, *η*_p_^2^ = 0.045), direction x speed (F(1,13) = 4.42, p=0.056, *η*_p_^2^ = 0.254) and task x direction x speed (F(1,13) = 0.14, p=0.713, *η*_p_^2^ = 0.011).

Together, these results suggest that MEPs differ depending on movement direction and speed, but not on visibility of the target curve. However, the direction effect as observed in the MEPs were also seen in the background EMG, which might have contaminated the MEP results.

#### Corticospinal excitability corrected for background EMG

To ensure that the effects of speed and direction for the MEPs were not due to differences in background EMG, we performed an additional analysis on the corrected MEPs. To correct for background differences in muscle contraction, we divided the MEP of each TMS event by the RMS100 in that event (shown in Figure 6B). For this MEP_EMG_ we found main effects of direction (F(1,13) = 12.5, p = 0.004, *η*_p_^2^ = 0.489) and speed (F(1,13) = 11.2, p = 0.005, *η*_p_^2^ = 0.464), indicating that MEPs were still higher when going up compared to going down (up = 0.29 ± 0.05; down = −0.22 ± 0.11) and moving fast compared to slow (fast = 0.10 ± 0.03; down = −0.03 ± 0.05), even when corrected for background EMG. For this analysis we also found an interaction for direction × task (F(1,13) = 4.8, p = 0.046, *η*_p_^2^ = 0.272). Post-hoc tests indicated that the upward movements were only different from the downward movements in the sinusoidal curve condition (up = 0.38 ± 0.07; down = −0.28 ± 0.04; p<0.001), but not in the horizontal line condition (up = 0.20 ± 0.09; down = −0.16 ± 0.13; p = 0.292). No differences were seen between the two conditions (between conditions: both p > 0.21). Finally, no significance for the main effect of task (F(1,13) = 0.16, p = 0.697, *η*_p_^2^ = 0.012), nor for the interaction effects of task x speed (F(1,13) = 2.22, p = 0.159, *η*_p_^2^ = 0.146), direction x speed (F(1,13) = 2.42, p = 0.144, *η*_p_^2^ = 0.157) and task x direction x speed (F(1,13) = 1.61, p = 0.226, *η*_p_^2^ = 0.110) was found.

#### Rate of force scaling

To assess whether participants performed the task correctly, we calculated the average speed prior to the TMS (Figure 6C). The ANOVA on the average speed showed significance for the main effects of direction (F(1,13) = 518.1, p < 0.004, *η*_p_^2^ = 0.98) and task (F(1,13) = 12.3, p = 0.004, *η*_p_^2^ = 0.486) but not of speed (F(1,13) = 0.2, p = 0.691, *η*_p_^2^ = 0.013). All interaction effects were significant: task x direction (F(1,13) = 7.0, p = 0.020, *η*_p_^2^ = 0.350), task x speed (F(1,13) = 41.9, p < 0.001, *η*_p_^2^ = 0.763), direction x speed (F(1,13) = 8.2, p < 0.001, *η*_p_^2^ = 0.877) and task x direction x speed (F(1,13) = 26.5, p < 0.001, *η*_p_^2^ = 0.671). Participants moved faster in the fast conditions compared to the slow conditions (all p<0.001), except in the horizontal line condition when making a downward movement where no difference was found (p = 1.00). Not surprisingly, the direction also differed based on whether up or down movements were made (all p<0.001). There were no notable differences between the sinusoidal curve and horizontal line conditions, except for the fast down condition, which was slower in the horizontal line compared to the sinusoidal curve condition (p = 0.002). These measures indicated that participants seemed to perform the task according to the instructions (sinusoidal curve: slow up: 0.83 ± 0.047, fast up: 1.35 ± 0.06, slow down: −0.73 ± 0.048, fast down: −1.60 ± 0.075; horizontal line slow up: 0.89 ± 0.039, fast up: 1.49 ± 0.039, slow down: −0.76 ± 0.085, fast down: −0.94 ± 0.169).

#### Motor performance

To determine whether participants’ performance differed between conditions, we measured the AUC of the difference between the target curve and the applied force (Figure 6D). An ANOVA on the AUC showed that the deviation depended on the direction (F(1,13) = 148.1, p<0.001, *η*_p_^2^ = 0.919), speed (F(1,13) = 161.7, p<0.001, *η*_p_^2^ = 0.926) and task (F(1,13) = 100.0, p<0.001, *η*_p_^2^ = 0.885). Participants deviated further from the curve when they moved upwards (up = 0.30 ± 0.01) compared to downwards (down = 0.21 ± 0.02). In addition, a larger AUC was seen for fast movements (fast = 0.32 ± 0.02) compared to slow movements (slow = 0.18 ± 0.01). Finally, larger deviations were seen in the horizontal line condition (mean = 0.33 ± 0.02) compared to the sinusoidal curve condition (mean = 0.17 ± 0.01). Furthermore, significant interactions for direction x task (F(1,13) = 56.0, p<0.001, *η*_p_^2^ = 0.812) and for speed x task (F(1,13) = 102.3, p<0.001, *η*_p_^2^ = 0.887) were observed. However, the post-hoc tests of these interactions were all significant (all p<0.01), providing no new interpretations to the data. Finally, no significance was found for the interaction effect task x direction x speed (F(1,13) = 1.4, p = 0.254, *η*_p_^2^ = 0.099)

In sum, CSE was primarily modulated by the direction and speed of the movement, even after correcting for background EMG. This CSE modulation was seen across in the sinusoidal curve and horizontal line conditions, i.e., when the sinusoidal target curve was visible or when only feedback from a straight line was present. As such, our findings mainly provide evidence that the motor system encodes ongoing motor components (i.e., direction and speed) but not that it encodes the differences in available information between the sinusoidal curve and horizontal line conditions.

### 5.2 Experiment 2

In experiment 2, participants lifted objects that were presented either in a series of increasing or random weights. After a series of increasing weights or within the random series, a ‘catch trial’ (weight 3) was presented. Immediately prior to this catch trial, in both the series of random and increasing weights, the heaviest (weight 5) was presented. In addition, we applied TMS prior to object lifting to assess CSE modulation during lift planning. With this experiment, we aimed to investigate whether lift planning (on the behavioral and corticospinal level) reflects implicit information, related to a series of random or increasing object weights, during a discrete object lifting task. For this we investigated differences in predictive force scaling and CSE modulation for the catch trial only.

#### Debriefing results

After the experimental task, participants were debriefed and were asked questions using a standardized questionnaire (see methods) related to the experimental task. All participants (Question 1: 14/14) reported that the weights of the object changed between trials. All but one participant (Question 2: 13/14) experienced these changes as random and two participants (Question 3: 2/14) reported to experience a pattern in the object presentation. While most participants (Question 4: 10/14) reported all weights to appear equally frequent, three answered that there were more heavy objects in the sequence and one participant argued they did not know whether the weights appeared equally frequent.

#### Corticospinal excitability

As can be seen in Figure 7A, neither the main effects of series (F(1,13) = 1.5, p = 0.705, *η*_p_^2^ = 0.011) and weight (F(1,13) = 0.65, p = 0.434, *η*_p_^2^ = 0.048) nor the interaction effect of series x weight (F(1,13) = 0.9, p = 0.373, *η*_p_^2^ = 0.061) were significant. As such, our results provide no evidence that objects weight influenced CSE modulation differently during the series of increasing and random weights. With specific interest to the series of increasing weights, no significant correlation was found between object weight and MEP amplitude (excluding catch trial: r = −0.134, p = 0.323; including catch trial: r = −0.115, p = 0.341).

#### Grip force at lift off

For GFatLO (shown in Figure 7B), the main effect of series (F(1,13) = 0.0, p = 0.851, *η*_p_^2^ = 0.003) and the interaction effect of series x weight were not significant (F(1,13) = 1.4, p = 0.258, *η*_p_^2^ = 0.097). In contrast, the main effect of weight (F(1,13) = 200.6, p < 0.001, *η*_p_^2^ = 0.939) was significant. Participants exerted more grip force at lift off when lifting the heavy weight (heavy = 16.40 ± 0.85 N) than when lifting the catch weight (catch = 12.16 ± 0.76 N). Notably, the correlation between object weight and the grip applied at lift off for the series of increasing weight was significant when both the catch trial was excluded (r = 0.655, p < 0.001) and included (r = 0.614, p < 0.001).

#### First peak grip force rate

For peak 1 GFR (Figure 7C), all effects were not significant: series F(1,13) = 0.0, p = 0.899, *η*_p_^2^ = 0.001), weight F(1,13) = 3.6, p = 0.078, *η*_p_^2^ = 0.220) and series x weight F(1,13) = 0.3, p =0.586, *η*_p_^2^ = 0.023). In addition, correlations between object weight and the first peak of grip force rate were not significant either (excluding catch weight: r = 0.067, p = 0.624; including catch weight: r = 0.059, p = 0.792).

#### First peak load force rate

For peak 1 LFR (Figure 7D), the effects of series F(1,13) = 1.3, p = 0.283, *η*_p_^2^ = 0.088) and series x weight F(1,13) = 1.5, p = 0.239, *η*_p_^2^ = 0.105) were not significant. However, the main effect of weight F(1,13) = 8.5, p = 0.012, *η*_p_^2^ = 0.395) was significant: participants scaled their load forces faster (larger peak LFR value) when lifting the catch weight (catch = 31.40 ± 2.43 N/s) than when lifting weight 5 (weight 5 = 28.64 ± 2.12 N/s). Akin to peak 1 GFR, correlations, correlations between object weight and the first peak of load force rate were not significant (excluding catch weight: r = 0.036, p = 0.791; including catch weight: r = 0.032, p = 0.792).

#### Loading phase duration

Similar to peak 1 LFR, the main effect of weight F(1,13) = 157.7, p < 0.001, *η*_p_^2^ = 0.924) was significant for LPD (Figure 7E) whereas the effects of series F(1,13) = 0.3, p = 0.592, *η*_p_^2^ = 0.023) and series x weight (F(1,13) = 1.7, p = 0.214, *η*_p_^2^ = 0.116) were not. In line with peak 1 LFR, participants had a shorter loading phase duration when lifting the catch weight (catch = 196.75 ± 9.28 ms) than when lifting weight 5 (weight 5 = 320.64 ± 14.47 ms). Notably, correlations between object weight and LPD were significant (excluding catch weight: r = 0.809, p < 0.001; including catch weight: r = 0.776, p < 0.001).

In sum, these findings show differences (Grip force at lift off and loading phase duration) on the behavioral level when participants lifted the heaviest weight or the catch one. However, our results provide no evidence that participants lifted the catch weight differently after the series of increasing or random weights. In addition, this absence of proof was also present on the corticospinal level (i.e., no differences between MEPs).

## 6. Discussion

In the present study, we investigated how explicit and implicit contextual information affected force scaling and CSE modulation during hand-object interactions. In experiment 1, participants performed a continuous force tracking task and in experiment 2, other participants performed a discrete object lifting task. In both experiments, motor performance was assessed to investigate how participants scale their fingertip forces during continuous and discrete hand-object interactions. TMS was used to probe the underlying changes in CSE.

For experiment 1, we found that when a sinusoidal curve was shown (i.e., explicit visual information of position on curve and performance error), participants performed significantly better compared to the condition in which the sinusoidal curve was shown as a horizontal line (i.e., explicit visual information of performance error but implicit information regarding position on curve). Our results indicate that CSE was modulated primarily by movement-related features (i.e., fast vs slow change in muscle contraction and up-vs downscaling of fingertip forces) in both the sinusoidal curve and horizontal line conditions. However, our findings do not robustly show that CSE modulation was altered based on how visual information was presented. As such, these findings indicate that CSE is primarily modulated by task requirements that can be directly mapped onto the corticospinal output (i.e., rate and direction of force scaling) but not by whether this information is explicitly or implicitly provided.

For experiment 2, we found no behavioral and neurophysiological differences when participants performed the catch trial after the series of increasing weights or after the series of random weights. These findings generally indicate that participants did not predictively scale their forces by extrapolating from the implicit weight increasing series.

In experiment 1, participants performed a sinusoidal force tracking task when provided with the actual sinusoidal target force (sinusoidal condition) or only received feedback about their performance in the form of a deviation from a horizontal line (horizontal line condition). As can be seen in Figure 6D, participants performed significantly worse (higher AUC value) in the horizontal line condition compared to the sinusoidal curve one. These findings indicate performance suffered when participants could not visually track their location on the sinusoidal curve and thus not predict the required directionality (up- or downscaling of force scaling) and rate of force scaling (the steepness of the curve’s slope). On the neural level we found that CSE modulation (Figure 6A-B) was primarily driven by ongoing movement-related features, i.e., direction and the change in force scaling rate (i.e., “speed”). CSE was larger when participants were upscaling their fingertip forces compared to downscaling (Figure 6A). The latter findings are in line with Kimura et al. (2003) and Li (2013) who showed that CSE is increased when contracting the elbow flexor (compared to relaxation) and when dynamically contracting the muscles of four fingers (without thumb) compared to isometric contraction and relaxation, respectively. These studies and our results agree with findings that motoneuron inputs having a greater relative contribution during shortening muscle contractions than relaxations (Semmler et al. 2002) and that CSE decreases prior to muscle relaxation (Suzuki et al. 2015). Importantly, our results extend on these studies by showing that (a) CSE is increased when upscaling fingertip forces compared to downscaling during a force tracking task, and (b) CSE modulation is also driven by movement speed, where it is increased with fast compared to slow changes in force.

Critically, effects of CSE modulation by the availability of explicit information (i.e., sinusoidal curve vs horizontal condition) seems to be minimal at best and most likely to be absent. As such, our findings indicate that CSE modulation reflected ongoing task execution (i.e., rate of force scaling and direction) but not the availability of explicit/implicit contextual information driving task performance.

Despite information regarding the task being less explicitly available in the horizontal line condition (i.e., error shown in inverse manner and no explicit information regarding the curve’s slope), CSE modulation was not driven differently than in the sinusoidal curve condition. Interestingly, these findings are in line with Young et al. (2000): using a reaction time task, they showed that when individuals respond to stimuli, CSE is automatically driven and not selectively modulated by intention- (‘readiness to respond’) and attention-selectivity (‘preferential processing of stimuli’). In addition, it has been proposed by Albert et al. (2022), using reaching tasks, that the amount of implicit learning can be altered by explicit strategies due to the implicit and explicit learning systems competing for errors as a common source of information about performance. Finally, Bestmann et al. (2008) showed that CSE modulation during action preparation becomes smaller when uncertainty increases. Combining the findings of these studies, it is feasible that, in our experiment 1, participants used the same behavioral strategy across both tracking conditions: focus on deviations between target line and actual performance and, thus, not necessarily on the target line’s shape. Given that errors were shown differently between conditions (i.e., inverse in the horizontal line condition leading to higher uncertainty), this could explain lower performance in the horizontal line condition while the similarity in the behavioral strategy between conditions could then explain the absence of differences in CSE modulation across conditions. However, this assumption cannot be verified with the current data but could be investigated in future studies including eye tracking to probe participants’ behavioral strategy and conditions where the visibility of the target line is manipulated. In sum, our findings for the continuous force tracking experiment suggest that corticospinal outputs are primarily driven by the ongoing motor commands and do not seem to be influenced by the visual nature of how feedback about ongoing performance is provided.

A potential limitation of our study is that we only performed TMS at the upper part of the curve, i.e., at 2.5 N. This could create a potential bias regarding the up and down conditions, since TMS was applied when participants were increasing their forces for a longer time compared to when they were decreasing the force. Although the forces were dynamic, requiring no adaptation to a static force, and the rate of change (frequency) could be altered at the 2N midway point, it is possible that participants adapted differently to specific force directions or force speeds. However, since our differences between upward and downward forces replicate results of previous studies (Kimura et al. 2003), it seems unlikely that differences in CSE modulation were only due to adaptation differences in behavior.

It should be noted that, since TMS was applied at a fixed force level, participants could have anticipated the TMS pulse. However, we took care to construct the sinusoidal curves in such a way that TMS was not always applied at 2.5 N: a minimum duration of 3.5 s was placed between each stimulation to avoid carry-over effects. As such, TMS was not applied every time participants exerted 2.5 N of force, thus making the stimulation less predictable. Finally, since we found no differences in CSE modulation between the horizontal and sinusoidal condition, it seem unlikely that the MEPs were affected by expectation of the TMS pulse. In support of the continuous motor task in experiment 1, we performed experiment 2 to investigate how implicit information affects CSE modulation in a discrete object lifting task. Previous studies have indicated that the CSE modulation represents task requirements (i.e., object weight) based on the previously lifted object (Loh et al., 2010). Noteworthy, behavioural evidence suggests that the sensorimotor system can adapt other object lifting strategies based on information acquired across several lifts (Mawase & Karniel 2010; Cashaback et al. 2017). Here, by relying on a paradigm similar to Mawase & Karniel (2010), we investigated whether a lifting strategy, acquired by lifting hidden series of increasing weights, would be reflected in CSE modulation. Participants performed an object lifting task on differently weighted, but visually identical, cubes. Within the overarching sequence of presented weights (see Figure 5), two types of series were hidden. One consisted of random weights whereas the second one consisted of increasing weights. Importantly, in both the random and increasing weights series, the second to last trial was always performed on the heaviest object and the last trial on a lighter ‘catch weight’, as in Mawase & Karniel (2010). The second to last weight was standardized across increasing and random series to control for sensorimotor memory effects (Baugh et al., 2012; Castiello, 2005; Johansson & Westling, 1993). With this experiment, we wanted to investigate whether CSE modulation would reflect the weight series (random or increasing) beyond each lift separately. Critically, our results provided no evidence that participants lifted the catch weight differently based on the series of increasing weights. Accordingly, we did not observe CSE modulation either. That is, we did not find statistical differences for behavioral parameters and CSE between same-weighted lifts after the series of random and increasing weights. Noteworthy, Mawase & Karniel (2010) reported that participants extrapolated their fingertip forces based on the hidden series of increasing weights. There are three major differences between their and our study that could explain why we could not replicate their behavioural results.

First, they relied on a virtual reality set-up whereas we used real hand-object interactions. In a recent review, Snow & Culham (2021) argued substantial differences between how virtual and real objects affect perception, cognition and action. For instance, individuals have a ‘real-object preference’ when gazing at real objects and 2D representations of the same object. It is possible that in the study of Mawase & Karniel (2010) participants relied on a virtual-reality associated lifting strategy that differs from how we interact with real-world objects. For instance, in a virtual reality setup, some tactile information (e.g., friction between object and fingertips) is missing, which contributes to force scaling (Castiello, 2005) thus potentially altering the grasping strategy. Second, as a proxy for predictive force scaling, Mawase & Karniel (2010) relied on grip force at lift-off whereas we relied on multiple dependent proxies (grip force, load force duration and force rates). It has been argued that the force rate provides a higher sensitivity for probing predictive lift planning (Johansson & Westling, 1993). Critically, neither for grip force at lift-off (akin Mawase & Karniel, 2010) nor for the other behavioral measures did we find differences between conditions. Third, Mawase & Karniel (2010) did not include a direct comparison between same-weighted lifts after the series of increasing and random weights. We argued that, if we wanted to investigate how predictive force scaling would be altered by the hidden series, a direct comparison between same weighted objects in different conditions should be made. This is the typical approach we used before (for example see: Rens & Davare, 2019; Rens et al., 2020; Van Polanen et al., 2020) and which has been used by other groups in our field as well (for example see Alaerts et al., 2010; Baugh et al., 2012; Buckingham et al., 2014; Cashaback et al., 2017; Gordon et al., 1993; Johansson & Westling, 1993; Loh et al., 2010; Reichelt et al., 2013; Senot et al., 2011). As such, it is unclear whether the results of Mawase & Karniel (2010) reflect (i) an object lifting strategy driven by implicitly learning the hidden series or (ii) solely sensorimotor effects that are entirely driven by last lifted object. However, it is evident in our experiment that participants did not develop a lifting strategy based on the hidden series. Potential explanations for why such strategies were not adopted by participants can be derived from relevant literature. First, Cashaback et al. (2017) showed that the sensorimotor system, in the case of uncertainty, aims to minimize prediction errors across many lifts instead of the prediction error for each lift separately. In our study, participants were instructed to perform the lifts naturally, but they were not specifically instructed to avoid lifting errors as in previous work from our group (Rens et al., 2021a). As such, participants might have relied on a general lifting strategy, minimizing prediction errors across all trials (akin to Cashaback et al., 2017) instead of attempting to “discover” the hidden series of increasing weights and predictively scale fingertip forces for each upcoming weight specifically (akin to Mawase & Karniel, 2010). Second, another explanation could be that the sensorimotor system prioritizes implicit information (akin to Albert et al., 2022) that is not related to the series of increasing weights. For instance, it is feasible that predictive force scaling, based on sensorimotor memories, relies on implicit motor learning (e.g., Flanagan et al., 2001). Apparently, single-lift information (i.e., sensorimotor memory) is more readily available than multi-lift information (i.e., series of increasing weights). As such, it seems feasible that the sensorimotor system prioritizes the integration of implicit information related to the previous lift but not to the entire series. Subsequently, if the participant explicitly understands the series to be random (and thus does not discover the implicit series of increasing weights), they might fall back on a general lifting strategy (again akin to Cashabak et al., 2017). Future research could tease apart some of the questions raised here. For instance, by altering the amount of explicit information (e.g., instructing to lift without lifting errors instead of lifting naturally or providing participants with distinct strategies) or implicit series of weights (e.g., having each presented multiple times in a row or use outlier weights for catch trials), it might be possible to further investigate how implicit information is integrated for discrete object lifting as has been done for reaching (Albert et al., 2022).

With respect to CSE, our results provide no evidence that CSE modulation was altered by the hidden series of increasing weights. Loh et al., (2010) showed that when individuals plan to lift an object, CSE was modulated by the previously lifted object. Based on Loh et al. (2010), it seems plausible that CSE modulation in experiment 2 depended entirely on the sensorimotor memory of the previous lift which reflected the heaviest weight that was the same in both the increasing and random series. It is unlikely that this null result is driven by the timing of the TMS pulse during lift planning as we did find CSE modulation in our previous work using the same TMS timing (Rens et al., 2021b). In addition, other work reported lift planning effects on CSE modulation when the TMS pulse was applied even longer after object presentation (Parikh et al., 2014). As such, and in support of our behavioral findings, we argue that the null results are caused by sensorimotor control mechanisms (i) not being able to detect the hidden series of increasing weights (ii) prioritizing other implicit information (Albert et al., 2022) or (iii) opting for a more general lifting strategy that is not trial-specific (Cashaback et al., 2017).

In conclusion, in the present study we investigated how corticospinal outputs could be modulated by implicit information during the execution of continuous (experiment 1) and discrete (experiment 2) hand-object interactions. Our results indicate that CSE modulation is primarily concerned with ongoing force scaling (experiment 1). Noteworthy, our results fail to demonstrate that contextual implicit information (experiment 1: type of visual feedback; experiment 2: hidden series of increasing weights) is represented into corticospinal outputs. As such, these findings indicate that sensorimotor control might not integrate implicit contextual information for hand-object interactions. Accordingly, other implicit information sources related to ongoing task performance during continuous force exertion and discrete object lifting might be prioritized.

## Acknowledgements

The authors would like to thank Immele Meeusen and Hannah De Buck for their help in the data collection of experiment 1 and 2, respectively. We also would like to thank René Clerckx for programming the force tracking task in experiment 1.

